# Modeling Progression of Single Cell Populations Through the Cell Cycle as a Sequence of Switches

**DOI:** 10.1101/2021.06.14.448414

**Authors:** Andrei Zinovyev, Michail Sadovsky, Laurence Calzone, Aziz Fouché, Clarice S. Groeneveld, Alexander Chervov, Emmanuel Barillot, Alexander N. Gorban

## Abstract

Cell cycle is the most fundamental biological process underlying the existence and propagation of life in time and space. It has been an object for mathematical modeling for long, with several alternative mechanistic modeling principles suggested, describing in more or less details the known molecular mechanisms. Recently, cell cycle has been investigated at single cell level in snapshots of unsynchronized cell populations, exploiting the new methods for transcriptomic and proteomic molecular profiling. This raises a need for simplified semi-phenomenological cell cycle models, in order to formalize the processes underlying the cell cycle, at a higher abstracted level. Here we suggest a modeling framework, recapitulating the most important properties of the cell cycle as a limit trajectory of a dynamical process characterized by several internal states with switches between them. In the simplest form, this leads to a limit cycle trajectory, composed by linear segments in logarithmic coordinates describing some extensive (depending on system size) cell properties. We prove a theorem connecting the effective embedding dimensionality of the cell cycle trajectory with the number of its linear segments. We also develop a simplified kinetic model with piecewise-constant kinetic rates describing the dynamics of lumps of genes involved in S-phase and G2/M phases. We show how the developed cell cycle models can be applied to analyze the available single cell datasets and simulate certain properties of the observed cell cycle trajectories. Based on our modeling, we can predict with good accuracy the cell line doubling time from the length of cell cycle trajectory.

## 1 Introduction

Cell cycle is the most fundamental biological process underlying the existence and propagation of life in time and space. Progression through the cell cycle represents a complex dynamical process, which involves coordinated changes at all levels of cellular organization, in particular, at transcriptomic and proteomic levels. The major players of the cell cycle machinery were characterized a long time ago, but its comprehensive description involving connection to other cellular subsystems is far from being completed [1, 2].

Progression through the cell cycle can be seen as a trajectory in multidimensional space of all possible cellular states, similarly to other processes such as differentiation or ageing. However, this trajectory has a particularity related to its periodicity. Simplifying biological reality, we can say that at the end of this trajectory a cell splits into two daughter cells twice as small. After this event, each of the daughter cells must find itself in a state corresponding to the starting point of the trajectory of its parent. This requirement imposes certain constraints on the geometry and underlying mechanisms of the cell cycle trajectory (CCT), which should be reproduced in any formal treatment of this process, such as mathematical modeling.

Cell cycle process has been a subject of mathematical modeling for many decades [3–5]. Most of the existing models focused on reproducing the logics of the regulatory interplay between protein actors, at the level of their expression and post-translational modifications. Multiple modeling formalisms have been exploited such as chemical kinetics [6, 7], logical modeling [8], Petri nets [9], approaches based on tropical algebra [10, 11]. The transcriptional dynamics of cell cycle have been much less addressed by mechanistic mathematical modeling so far.

Introducing high-throughput omics measurements gave rise to a number of molecular studies with the objective to characterize each cell cycle phase in terms of associated molecular changes such as sets of specifically expressed genes [1, 12]. The interest in the molecular organization of cell cycle has been boosted after the appearance of single cell technologies, which allowed us to visualize and study the cell cycle trajectory explicitly without synchronizing individual cells which can be problematic, especially in vivo. Recent single cell transcriptomic and proteomic studies allowed molecular description of progression through the cell cycle in continuous fashion. Such representation not only simply delineates the cell cycle phase borders but also characterizes each cell for its precise progression position within each phase [13–16].

Better understanding cell cycle functioning is of utmost importance for cancer research, where the deviation from the normal cell cycle progression is expected. One has to be able to answer the following questions: What is the normal pattern of the events comprising a cell cycle, and to what extent does it vary in normal physiology? What deviations from a normal cell cycle are characteristic for a tumor cell? What processes trigger these changes and are they specific to a cancer type? and many others. Mathematical modeling represents a tool for rationalizing and formalizing reasoning on these complex questions.

Application of existing mathematical models of cell cycle to answer these questions remains difficult because of discrepancy between the nature of the data and the level of the description present in the models. Thus, the most comprehensive data source we currently possess is at the level of transcriptomic changes in single cells, while the existing modeling efforts focus on protein players. The data shows involvement of hundreds of genes and proteins in cell cycle dynamics, while the models include a tiny fraction of this number manipulating with few tens of players at most. Therefore, we believe that the development of mathematical models matching the complexity and the nature of the abundantly available data is still highly needed. In particular, we believe that even a simple mechanistic model of cell cycle transcriptome dynamics, capturing its main features, is lacking in the field. It appears that using dynamical variables representing relatively large lumps of genes (e.g., all genes involved in DNA replication) might be a useful coarse-grained approach to modeling cellular transcriptomes. The motivation of this study is to contribute to closing this gap.

Single cell studies of cell cycle trajectory in snapshots of actively proliferating cells represent a unique opportunity to formulate the most general principles underlying such modeling. In [17] the principle of minimizing transcriptomic acceleration has been recently suggested. According to this principle, the transcriptomic cell cycle trajectory should represent a spiral, or, after neglecting the relatively slow drift unrelated to cell cycle progression, a shape close to “a flat circle”. The trajectory of approximately this shape was indeed phenomenologically observed in the HeLa cell line profiled with scRNASeq technology, after deconvoluting the transcriptomic dynamics connected to the cell cycle from other sources of transcriptional heterogeneity. In particular, the absence of cell cycle-related transcriptional epochs was deduced from this model.

In the current study, we suggest an extended and different point of view on the properties of transcriptomic cell cycle trajectory which, in our opinion, in some cases better matches its observed properties in various cellular systems, when sufficiently good quality data can be collected. We propose a formal model of CCT as a sequence of epochs of growth during each of which the trajectory is approximately linear in the space of logarithmic coordinates. Therefore, CCT can be modeled as a piecewise linear trajectory in the space of logarithms of some extensive cell properties, followed by a shift at the vector with coordinates – log 2 which represents the cell division event. This model explicitly assumes existence of well-defined transcriptional epochs in CCT.

Movement along linear trajectory in the space of logarithms of the values of some cellular properties means that along the trajectory any two such properties *x_i_, x_j_* are connected through a power law dependence 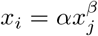, *α, β* = const. Such dependencies are known as allometric in many fields of biology [18–22]. Some approaches in mathematical chemistry and theoretical biology, dealing with systems in stable non-equilibrium, exploit the linear relations between chemical potentials which can be expressed as logarithms of species concentrations [23, 24].

Particular cases of allometric dependencies are when all the quantities grow linearly with physical time, or when all the quantities follow exponential growth or decay *x_i_* = *b_i_* exp(*a_i_t*). The model of movement along piecewise linear trajectories with an event of cell division represents the simplest scenario which is easy to analyse theoretically and to simulate. Nevertheless, the most important conclusions derived from this analysis will stay valid for the trajectories that do not deviate too much from linearity which is the case for available experimental data.

Using the model of piecewise linear growth with division, we formulate a fundamental statement about correspondence between the number of linear segments in the cell cycle trajectory *m* (number of the most important states of the cell cycle-related transcriptional machinery) and its effective embedding dimension *n*. First part of this statement (*m* ≥ *n*) has the character of a strict theorem with formal proof. The second part (*m* ≤ *n*) has a character of a feasible hypothesis which can be validated using available data. The statement *m* = *n* suggests that the transcriptomic cell cycle trajectory is essentially “not flat”, since the number of segments we can observe can be as high as 4 or 5.

The type of model that we discuss here was partly introduced in [25] by a group of authors standing behind the collective pseudonym *Shkolnik*, with several authors of the current manuscript belonging to the group. Here we revise the notations of the model, significantly extend it and adapt to describing the cell cycle trajectory in single cell datasets which became recently available.

In order to connect the geometric properties of cell cycle trajectory to interpretable mechanistic parameters, we extended the model of piecewise linear growth in logarithmic coordinates, to a simple kinetic model with rates depending on time as piecewise constant functions. In this case, some of the segments of the trajectory become nonlinear but remain smooth and do not deviate from linearity too far. We used this model to obtain estimates of the transcriptional epoch durations in physical time, and to simulate the changes in the CCT geometry as a result of these parameter modifications. We demonstrate that in some cases, the geometry of CCT can significantly deviate from the “simple circle” shape. Moreover, we formulate a quantitative prediction of the cell line doubling time, based on estimating the length of the cell cycle trajectory.

## 2 Results

### 2.1 Example of a cell cycle trajectory extracted from single cell data

The current study is motivated by the observation that after appropriate pre-processing of single cell RNA-Seq data (see Methods), one can observe the cell cycle trajectory (Figure 1) which can be approximated by piecewise linear curve, with a gap between the beginning and the end of the trajectory corresponding to the cell division moment.

**Figure 1:**
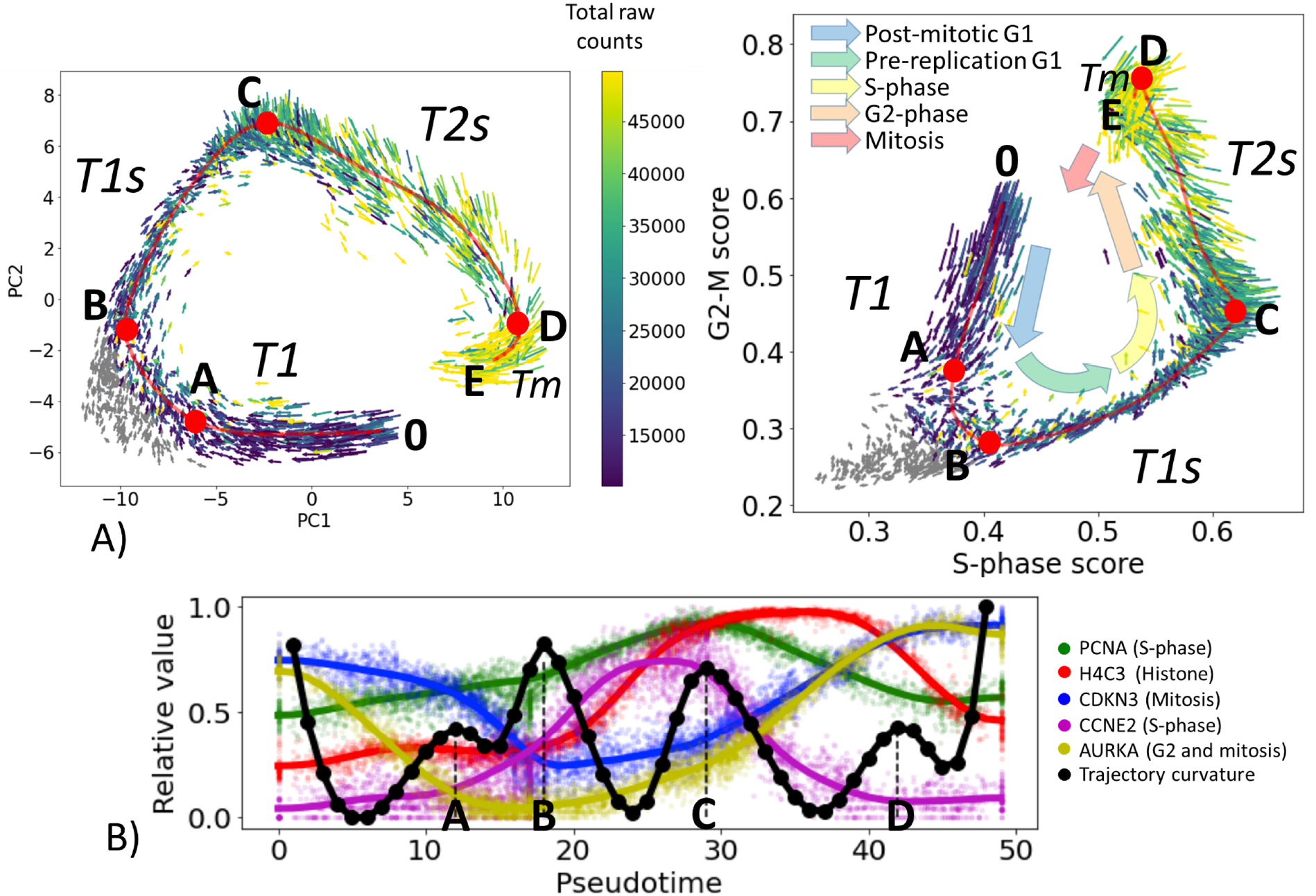
Cell cycle trajectory (CCT) of CHLA9 Ewing sarcoma cell line in the single cell transcriptomic space. A) Each cell is represented by an arrow reflecting the momentary direction and the speed of transcriptomic changes, estimated with RNA velocity. Two projections are shown, in the first two principal components and in the plane of S-phase and G2-M scores. The color of the arrows signifies either the total amount of RNA counts in the single cell profile (blue to yellow scale) or the cells in non-proliferative state (shown in grey). Red line shows an approximation of the cell cycle trajectory with a principal curve computed with ElPiGraph, directly in the 30-dimensional space of the first principal components of the dataset. Several particular positions along the trajectory (A,B,C,D) mark either the peaks of the Riemannian curvature of the principal curve (also shown in B) panel) or the beginning (0) and the end (E) of the trajectory. B) Pseudotemporal transcriptomic dynamics of several cell cycle-related genes along CCT, shown relatively to the maximum value units. The pseudotime range is from 0 to 49, corresponding to the number of nodes in the approximation of the principal curve (50 nodes). In black, an estimation of the Riemannian curvature of the principal curve is shown, with peaks indicated by letters (A,B,C,D).

Here we use the example of Ewing sarcoma cell line CHLA9 sequenced at single cell level using the Chromium 10x technology [26]. The particularity of this dataset was in that it contained a significant number of proliferating cells with single cell transcriptomes of good quality (more than 4000 cells with the total number of Unique Molecular Identifiers (UMIs) between 10000 and 50000). Also, the proliferation signal in this dataset seems to explain the largest fraction of transcriptomic heterogeneity, since in the plane of the first two principal components one can observe clear visualization of the cyclic trajectory. In other cell line single cell datasets, the proliferative signal can be masked by other sources of transcriptomic heterogeneity, requiring special procedures of data treatment to reveal it [17, 27, 28].

The scRNA-Seq data have been normalized in order to preserve the pattern of dynamics of the total number of counts (UMIs) along the CCT, see Methods section. The normalized gene expression levels are represented at the logarithmic scale, following the standard practice. Then multi-dimensional distribution of single cell transcriptomic profiles projected into the space of the first 30 principal components has been approximated by a principal curve, using the approach described in Methods. The curvature of the principal curve has been estimated using the standard formulas from the differential geometry, which revealed the existence curvature peaks, reflecting the rapid turning points of the trajectory. We hypothesized that these turning points correspond to the large-scale changes in the transcriptional programs of the cell cycle process. The pattern of momentary velocities of the transcriptomic changes, estimated with RNA velocity, was compatible with this hypothesis (Figure 1,A).

The pseudo-temporal dynamics of the known cell cycle-related genes confirmed that the trajectory curvature peaks delineates biologically meaningful transcriptional epochs. Thus, the epoch 0-A-B can be understood as a part of the G1 phase of the cell cycle, B-C as significantly overlapping with G1- and S-phases, and C-D as overlapping with S- and G2-phases. The epoch D-E can presumably reflect the relatively short M phase (mitosis). Accordingly, we denote the identified transcriptional epochs as T1, T1s, T2s and Tm. We should note that due to the delay between the gene and protein expression, the transcriptional epochs identified here can not correspond to the exact durations and the borders of the cell cycle phases.

Remarkably, within each of the identified transcriptional cell cycle epochs, the global dynamics of the transcriptome remain close to linear in the logarithmic scale. This allows us to suggest a simple model which can, for example, represent the collective dynamics of the genes related to the S-phase and G2/M phases (see below).

### 2.2 Model of cell cycle as a trajectory of allometric growth with switches and divisions

Based on the observations of the properties of cell cycle trajectory in several scRNASeq datasets, we hypothesized that it can be recapitulated by a formal model of linear growth in logarithmic coordinates with switches and a cell division event.

Let the state of a proliferating cell be determined by *n* substances quantified in their amounts, not in concentrations. Instead of their natural units (such as RNA counts), let us use the logarithms of these amounts. The cell is represented as an *n*-dimensional vector, and all possible combinations of these vector components define the cell configuration space. For our model it is important that the considered *n* quantities are extensive measures, not intensive ones; extensiveness here means that the total amount of a substance is a sum of the amounts found in different parts of a cell. A division (for two almost equal) “younger” cells is formalized as a shift by the vector with all components equal – log 2 in this space. A relevant example of extensive quantity is the total amount of RNA molecules present in a cell, or the amount of any specific subset of RNA molecules, i. e. representing mRNAs of the genes involved in a particular process (such as mitosis or S-phase).

We also assume that the cell is characterized by one of several (few) internal states. In each of these states, the cell finds itself within an epoch which is characterized by a linear trajectory in the cell state space. This trajectory extends until the cell will meet a condition, where a switch into another internal state of the cell happens, which changes the direction of the trajectory. For simplicity, we assume the conditions of a switch can be described by a linear function. The cell movement continues until a particular condition will be met in which the cell division event is triggered leading to the aforementioned translation of the vector representing the cell state.

Let us introduce some mathematical notations and consider a deterministic automaton *A* whose complete state is represented by a pair (*x, s*), where *x* ∈ *R^n^* is a vector in n-dimensional space, and *s* ∈ *S* is an integer number from a finite set *S* = {*S*_1_,.., *S_m_*}. In further we will call x a position of *A* and *s* an internal state of *A*. We will denote the automaton *A* in position *x* and in the internal state *s* as *A*(*x*|*s*).

Each internal state *S_k_, k* = 1..*m* is parametrized by a vector *a_k_* ∈ *R^n^* and, possibly, by a linear manifold *D_k_* of dimensionality *n* – 1 embedded in *R^n^* (hyperplane), which we will call “the cell division hyperplane”. *D_k_* can be undefined, in this case we denote *D_k_* = *null*.

Let us also introduce a set of *p* functions *G* = {*g*_1_,…, *g_p_*}, *g_i_*: *S* → *S*, which we will call switches. Each switch *g_i_* is a map which converts an internal state *s_j_* ∈ *S* into another internal state *s_r_* ∈ *S*. Each switch *g_i_* is parametrized by a hyperplane *L_i_* existing in *R^n^* and inducing the switch function *g_i_* each time the trajectory of the automaton intersects *L_i_* (see Figure 2,A).

**Figure 2:**
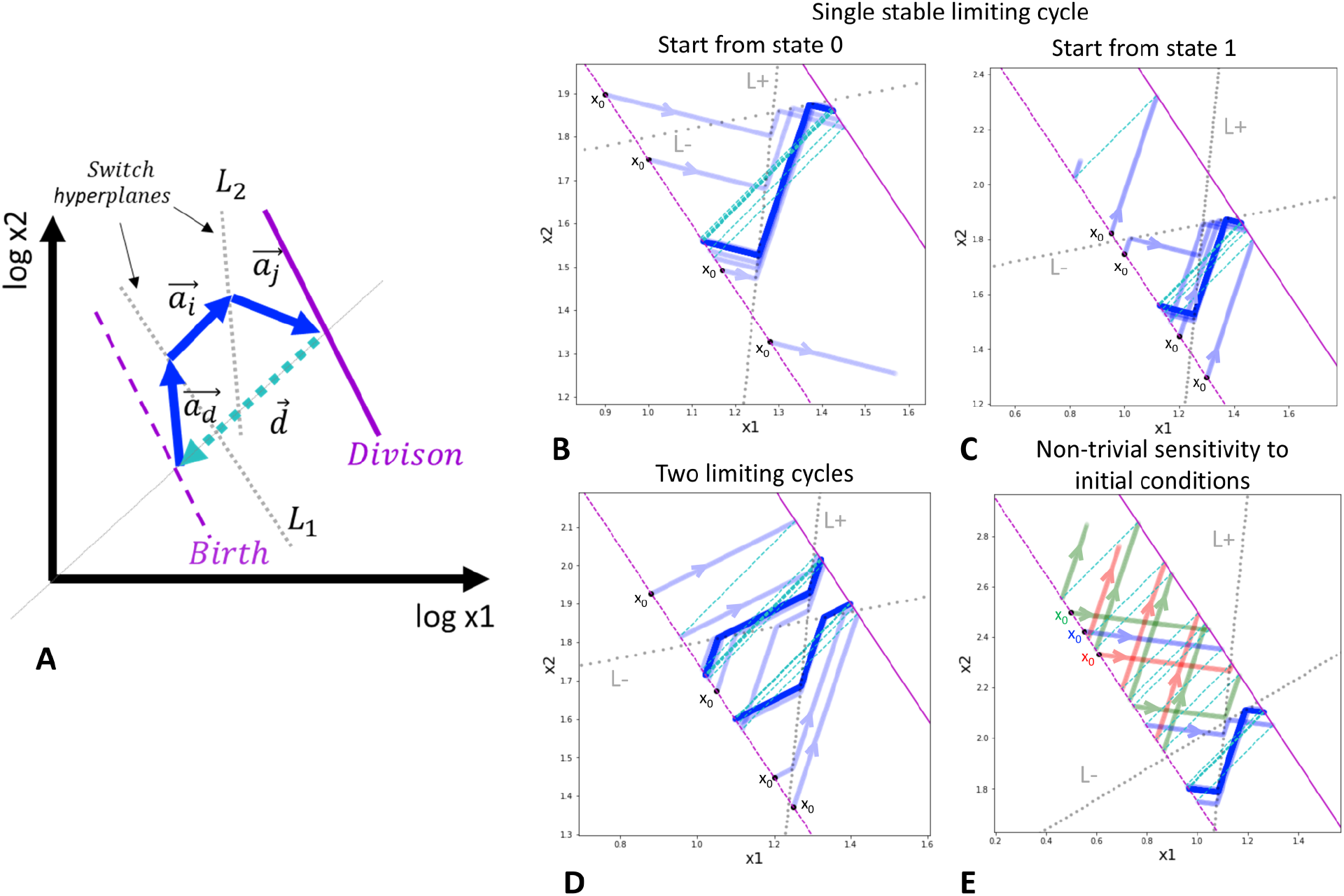
General schema of switch-like dynamics (A) and a toy model with a single trigger (B-E). A) Imaginary two-dimensional example of a limiting trajectory with division. The division hyperplane *D* is shown in purple, solid line. The birth hyperplane *B* is obtained from *D* by translation at vector *d*, shown in cyan (the most natural is to assume all the components of *d* to be – log 2). Two switch hyperplanes *L*_1_ and *L*_2_ are shown by grey lines. The limiting cycling trajectory is shown by blue arrows. B) and C) Example of single limiting cycle in the switching dynamics. Depending on the initial state of the automaton and the initial position, the trajectory enters into the limit cycle or degenerates (goes to infinity). For the same parameters, four initial conditions are shown. The trajectory is plotted with semi-transparent blue color such that the intense blue line designates the trajectory cycling multiple times on top of itself. D) Possibility of existence of two limit cycles. Depending on the initial state and position, the automaton ends up in one of the two possible limit cycles. E) Example of non-trivial dependence of the switching dynamics on the initial position of the automaton. The trajectories drawn by different colors from three closely located initial positions are shown, with two leading to degenerated dynamics and one located in between the first two, leading to the limit cycle. In B)-E) panels, the initial position of the automaton is always shown at the birth hyperplane *B* (shown my dashed purple line), therefore, it is characterized by a single number.

Finally, we introduce the cell division event *ϕ* which is a map between two states of *A*, such that *ϕ*((*x, s*)) → (*x* + *d, s_d_*), where *d* ∈ *R*^*n*−^ is a vector with negative components, and *s_d_* ∈ *S* is one of the possible internal states of *A*.

We will characterize any hyperplane here by a linear functional *f*(*x*|*b, c*) = *b*+ < *c, x* >, *b* ∈ *R, c* ∈ *R^n^*, where <, > denotes the standard scalar product between two vectors. Using such a functional, for any pair of vectors *x_i_, x_j_* ∈ *R^n^* we can determine if the linear segment connecting *x_i_* and *x_j_* intersects the hyperplane or not. If the segment intersects the hyperplane then *f*(*x_i_*) *f*(*x_j_*) < 0, and if it does not intersect then *f*(*x_i_*) *f*(*x_j_*) > 0. *f*(*x_i_*) *f*(*x_j_*) = 0 is satisfied only in a non-general position when either *x_i_* or *x_j_* is located exactly on the hyperplane.

The update rules for the automaton *A* are described as follows. The automaton starts in some initial position *x*_0_ and the internal state *s*_0_, and starts to move along the linear trajectory described by the equation *x* = *x*_0_ + *a*_0_*t*, where *a*_0_ is the vector of movement associated with the state *s*_0_. This movement continues unless one of the two events happen. In the first case, *A* reaches the corresponding cell division plane *D*_0_ (in case *D*_0_ is not null). Then the cell division event is triggered, *A*(*x*|*s*) → *A*(*x* + *d*|*s_d_*). In the second case, *x* reaches a switch hyperplane *L_j_* and then a switch of the internal state of *A* happens without changing its position, *A*(*x*|*s*_0_) → *A*(*x*|*g_j_*(*s*_0_)). The movement continues along a new trajectory, corresponding to the new cell state, following the same rules: either the trajectory hits the cell division hyperplane or any of the switch planes.

To summarize, the automaton *A* is characterized by its position and the internal state, see Figure 2,A. The asymptotic (in the infinite time limit) temporal dynamics of *A* is parametrized by a set of cell division planes *D* = *D_i_, i* = 1…*k*, set of switch functions *G* = {*g_i_*}, *i* = 1…*p* and the corresponding switch hyperplances *L* = {*L_i_*}, *i* = 1…*p*, and the parameters of the cell division event (namely, the translation vector *d* and the state after cell division *s_d_*).

It is convenient to encode the state *s* as a binary sequence of length *r* representing the on-off states of *r* triggers. In this case, a switch can be thought of as changing only one particular trigger from on to off or vice versa. In many situations, this makes the description of switch functions *g*: *S* → *S* quite natural as it will be seen below. Also, the state of the trigger might be not strictly binary but be characterized by several discrete positions, for example {0,1, 2}, just as it is the case in modeling multi-level discrete dynamics, where each discrete variable can take a value from a pre-defined finite set of levels.

The exact asymptotic trajectory of the automaton *A* can, in principle, depend on the initial position *x*_0_ and the initial internal state *s*_0_ of *A*.

During the temporal dynamics, some of the quantities can theoretically start moving towards negative or positive infinity. In the case of negative infinity, this means that the corresponding cellular quantity approaches zero. If not all quantities approach zero in the infinite time limit then we can say that in the stationary cell cycle, the quantity with zero value is not needed, and we can exclude it in the very beginning from the analysis. If all quantities go to zero in the infinite time limit then we can say that the functioning of the cell cycle is not possible (degenerates) with these parameters. In the case of growth of some of the quantities towards the positive infinity, we can assume that the description of the cell cycle progression is incomplete and some important switches are missing.

### 2.3 Simple example of dynamics with switches and cell division events

In the above-described switch-like dynamics, one can find examples of relatively complex behaviours even for the simplest scenarios, see Figure 2,B-E. Let us make a demonstration of this fact assuming that the dividing automaton is characterized by a position vector *x* with only two coordinates *x*_1_, *x*_2_, and its state *s* is encoded by only one binary trigger, so the automaton can be in two states *s* = 0 and *s* = 1, characterized by two vectors of movement *a*_0_ and *a*_1_, correspondingly. In order to be able to modify the trigger in both directions, we have to introduce two switch hyperplanes *L*^(+)^ and *L*^(−)^ with corresponding switch functions *g*^(+)^ = 1 (switch trigger on) and *g*^(−)^ = 0 (switch trigger off). Note that in this case the switch functions are constant, i.e. they map any state (which can be either 0 or 1) to a particular state. Let us also assume that the division event changes the automaton position but does not change its internal state.

In this simple toy example, by slightly varying parameters of the switching hyperplanes and the movement vectors, one can observe several interesting scenarii. In the most basic one either convergence of the trajectory to the limit cycle or divergence to infinity, indicating inconsistency of the chosen parameters with switching dynamics, is observed dependent on the initial state and the position of the automaton on the birth hyperplane, Figure 2,B-C. The switching dynamics trajectory can be also characterized by two limit cycles that can be achieved from different initial states and positions (Figure 2,D). Interestingly, when the initial position of the automaton is chosen quite far from the limit cycle, one can observe an effect of non-trivial sensitivity to the initial conditions (Figure 2,E). More precisely, the birth hyperplane can be split into a sequence of alternating intervals of equal length such that starting from one interval, the dynamics finally converges to the limit cycle, and starting from another interval, the dynamics diverges to infinity.

### 2.4 Two-dimensional model of cell cycle progression, fitted to the single cell transcriptomic data

Let us denote the aggregate signal related to the activation of genes associated with the S-phase of the cell cycle program as *S*, and the signal related to the activity of genes in G2 and *M* phases as *M*. Therefore, we will characterize the position of the automaton by a vector (*x_S_, x_M_*), just as it is presented in Figure 1,A, right panel. Let us denote the position of the turning points in the trajectory as (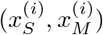, where *i* ∈ {0, *A, B, C, D, E*}.

We will encode the state of the system by the levels of two triggers, one associated with the S signal and another associated with the M signal. The three levels are denoted as a set {2 = *synthesis*, 1 = *decay*, 0 = *degradation*}. Intuitively, these levels correspond to the state of active transcription of the corresponding set of transcripts (‘synthesis’), absence of active transcription in which the transcripts are passively degraded accordingly to some base rate, and the process of active degradation when the transcripts are degraded more rapidly than the base rate. The state of the system is thus encoded by a pair of 3-level numbers *i, j* ∈ {0, 1, 2}. Accordingly, the 2D vectors of linear movement *a_ij_* are encoded by six rates 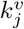, *i* ∈ {0, 1, 2}, *v* ∈ {*s, m*}, such that 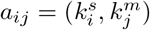. Accordingly to the intuition behind the introduced trigger levels, we assume constraints 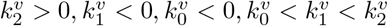.

Let us introduce 3 switches. First switch *g*_1_ turns on the synthesis of both variables independently on the current state, i.e. *g*_1_: (•, •) → (2,2), where • designates any level of the trigger. The second switch turns off the synthesis of genes in S-phase: *g*_2_: (2, •) → (1, •). The third switch turns off all the transcription independently on the current state, *g*_2_: (•, •) → (1,1). We assume that the division is possible only in the state (1,1) with transcription switched off, and that after the division event, the cell enters into the state of active degradation of the cell cycle genes (0,0).

The three introduced switches will be characterized by the corresponding switching hyperplanes. The first switch is triggered when the sum of the collective aggregated levels of expression of the genes involved in S and G2/M phases will reach some minimum *c_min_*, therefore, the linear functional associated with the first switch hyperplane is *f*_1_(*x_s_, x_m_*) = *x_m_* + *x_s_ – c_min_*. The second switch is triggered whenever the collective aggregated level of expression of S phase-associated genes will reach some maximum value *S_max_*, therefore, the linear functional associated with the second switch hyperplane is *f*_2_ (*x_S_, x_M_*) = *x_S_* – *S_max_*. Finally, the third switch is triggered when the collective aggregated level of expression of G2/M phase-associated genes will reach some maximum value *M_max_*, therefore, the linear functional associated with the third switch hyperplane is *f*_3_(*x_S_, x_M_*) = *x_M_ – M_max_*.

In the end, the cell division event will be triggered when the collective aggregated level of expression of G2/M phase-associated genes will cross some threshold *M_e_*, therefore, the linear functional associated with the division event is *f_d_*(*x_S_, x_M_*) = *M_e_* – *x_M_*.

Let us define the number of parameters in this simple switching model. Three introduced switches are characterized by 4 parameters *c_min_, S_max_, M_max_, M_e_*. There exists 6 rates *k_i^v^_* characterizing the movement vectors in the 9 = 3^2^ possible states, corresponding to all possible combinations of trigger levels. However, qualitatively, the dynamics in each automaton state is determined only by the direction of the corresponding vector and not it’s amplitude: therefore, we need one parameter per state visited during the progression through the cell cycle. Under certain constraints on the rates formulated above, and also on the switch parameters (namely, *c_min_ < S_max_, M_max_, M_e_ < M_max_*), the suggested model is constructed such that along the cell cycle trajectory only 4 states will be visited in a pre-defined order: (0,0) → (2,2) → (1,2) → (1,1). Therefore, the total number of parameters equals 8.

Knowing the position of four characteristic points along the cell cycle trajectory, namely 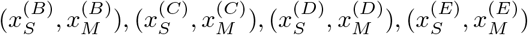, it is possible to completely parametrize the automaton. The starting and the end point of the cell cycle trajectory must be connected by the relation 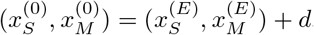, where *d* is the vector with components (– log_10_ 2, – log_10_ 2).

Therefore, we put 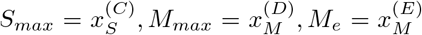. Instead of using directly the *B* point, we will use the position of the non-proliferating cell with the maximum sum of the coordinates in the *S, M* plane, and we designate it as 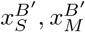 (other choices are also possible). Then 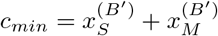. Then we define rates:

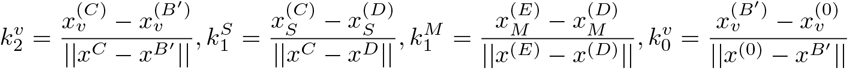

The resulting steady state cell cycle trajectory is shown in Figure 3.

**Figure 3:**
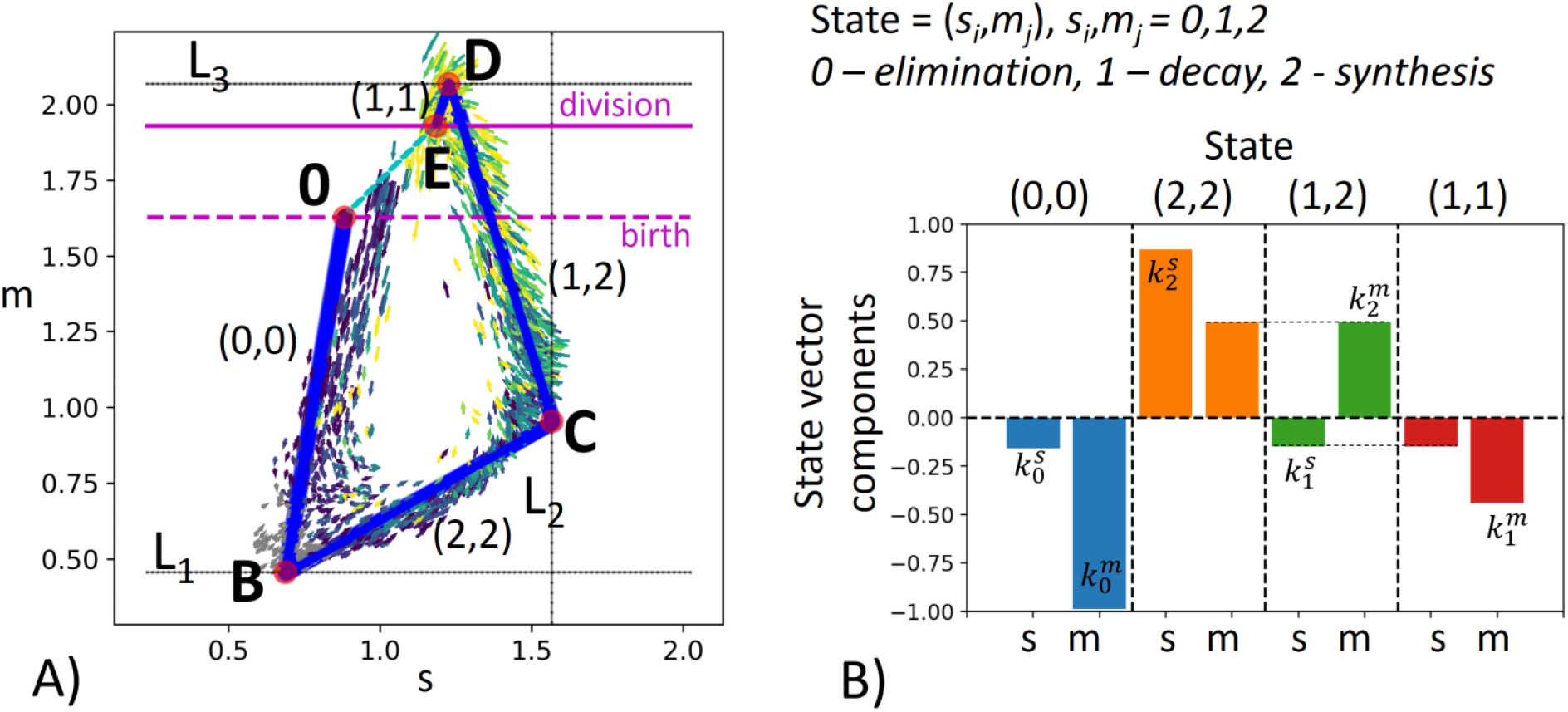
Modeling transcriptomic cell cycle trajectory by an allometric growth with switches. A) Piecewise linear cell cycle trajectory fit to the single cell RNASeq data (cell cycle trajectory, shown in Figure 1,A, right). The model contains three switching planes *L*_1_, *L*_2_, *L*_3_, and is characterized by 4 states. The states are encoded with two triggers, each possessing three possible levels 0,1,2, the biological meaning of which is specified in B). B) The growth vectors associated with each state are encoded by rates 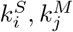, such that the components of the growth vectors equal 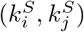, where *i* and *j* are the levels of the corresponding triggers.

We denote the linear segments of the trajectory shown in Figure 3 as T1, Ts, T2, Tm, assuming that they have significant overlap with G1, S, G2 and M phases correspondingly.

The suggested model describes 2D dynamics of the signals *S, M* which are empirically shown to explain most of the variance of all cell cycle genes in scRNASeq data (see below). However, higher-dimensional generalization of the suggested model is always possible. Also, in the model, we simplified the observed dynamics in Figure 1,A, left which seems to contain 5 segments, with an additional curvature peak in point A. The segment A-B seems to contain non-proliferating cells, and might correspond to the transcriptional epoch most similar to the quiescent cell state, when the active degradation of the mitotic transcripts was completely finalized. The existence of this epoch is less pronounced in the *S, M* projection (Figure 1,A, right), therefore we merged segments 0-A and A-B’ as the first order approximation.

### 2.5 Connection between the effective embedding dimensionality of cell cycle trajectory and the number of internal states

The introduced cell cycle modeling framework is a simple and empirical model, lacking mechanistic details. It’s main advantage is the possibility of analytical analysis of the most general necessary properties of the cell cycle trajectory geometry.

This geometry is embedded into a space of omics measurements, which dimensionality might be huge (e.g., expression of thousands of genes). However, we can assume that the intrinsic dimensionality (ID) of CCT is much smaller and that the state of the cell cycle progression can be characterized by *n* extensive variables, where *n* is relatively small. We will refer to *n* as CCT embedding dimensionality. Empirically it can be estimated by studying the snapshot of dividing single cells profiled with a particular technology, and computing its global intrinsic dimensionality (ID), provided that we could get rid of other non cell cycle-related sources of heterogeneity in measurements. Estimating ID can be done using one of the many existing methods for ID estimation [29, 30].

Let us establish the expected relation between *n* and the number of internal states *m* of the automaton approximating CCT. We intend to claim that theoretically *n* should match m under some natural assumptions.

We first state that *m* can not be smaller than *n*. In the theory of allometric growth with switches this statement has a character of strict theorem (see below), *m* ≥ *n*. Secondly, we state that *n* is expected to be at least equal to *m*. Both statements are of “general position” consideration type. However, the former one is strictly necessary, while the latter one represents a feasible hypothesis.

#### Theorem on the number of intrinsic cell cycle states

The number of segments m in the cell cycle trajectory modeled by the automaton with switches and linear growth in logarithmic coordinates must be not less than the cell cycle trajectory intrinsic dimensionality n, or m ≥ n

*Proof*. Let us consider the CCT dynamics in its *n* intrinsic coordinates each of which represents an extensive variable. The variable extensiveness means, in particular, that its value, after the cell division moment, divides by two. In logarithmic scale the cell division corresponds to the shift by vector *d* ∈ *R^n^* with n coordinates each of which equals – log 2. Each internal state is associated with a growth vector *a_i_* ∈ *R^n^, i* = 1..*m*. All non-negative linear combinations of *a_i_* form a convex cone 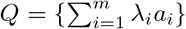, *λ_i_* ≥ 0. If *m* ό *n* then the set of vectors {*d*, {*a_i_, i* = 1..*m*}} is almost always linear independent and −*d* ∉ *Q*. Hence, −*d* is linearly separable from *Q*, according to the standard separability theorems. Linear separability of a point from a convex cone can be expressed as that for any non-zero *x* ∈ *Q* we can find a linear function *l*() such that *l*(*d*) = 0 and *l*(*x*) > 0. This makes the periodic cell cycle model impossible, because function *l*(*x*) increases along any growth direction, since for any *i* and *λ* > 0 we have *l*(*x* + *λa_i_*) = *l*(*x*) + *λl*(*a_i_*) > *l*(*x*), and after cell division *l*() does not change since *l*(*x* + *d*) = *l*(*x*) + *l*(*d*) = *l*(*x*). Therefore, the necessary condition of existence of stable cell cycle trajectory is *m* ≥ *n*, when the set of vectors {*d*, {*a_i_, i* = 1..*m*}} is linearly dependent, and also such choice of *a_i_* that −*d* ∈ *Q*. Only in this case one can satisfy the cyclic condition 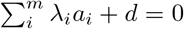 in general position of vectors {*d*, {*a_i_, i* = 1..*m*}}.

In simple words, this means that if *m* < *n* then in a general position the act of each cell division (shift by *d*) moves a cell state out of the subspace defined by the growth vectors. The only way to make the trajectory stay in this subspace is to make the cell division vector d belong to this subspace that can be guaranteed only if *m* ≥ *n*, see Figure 4.

**Figure 4:**
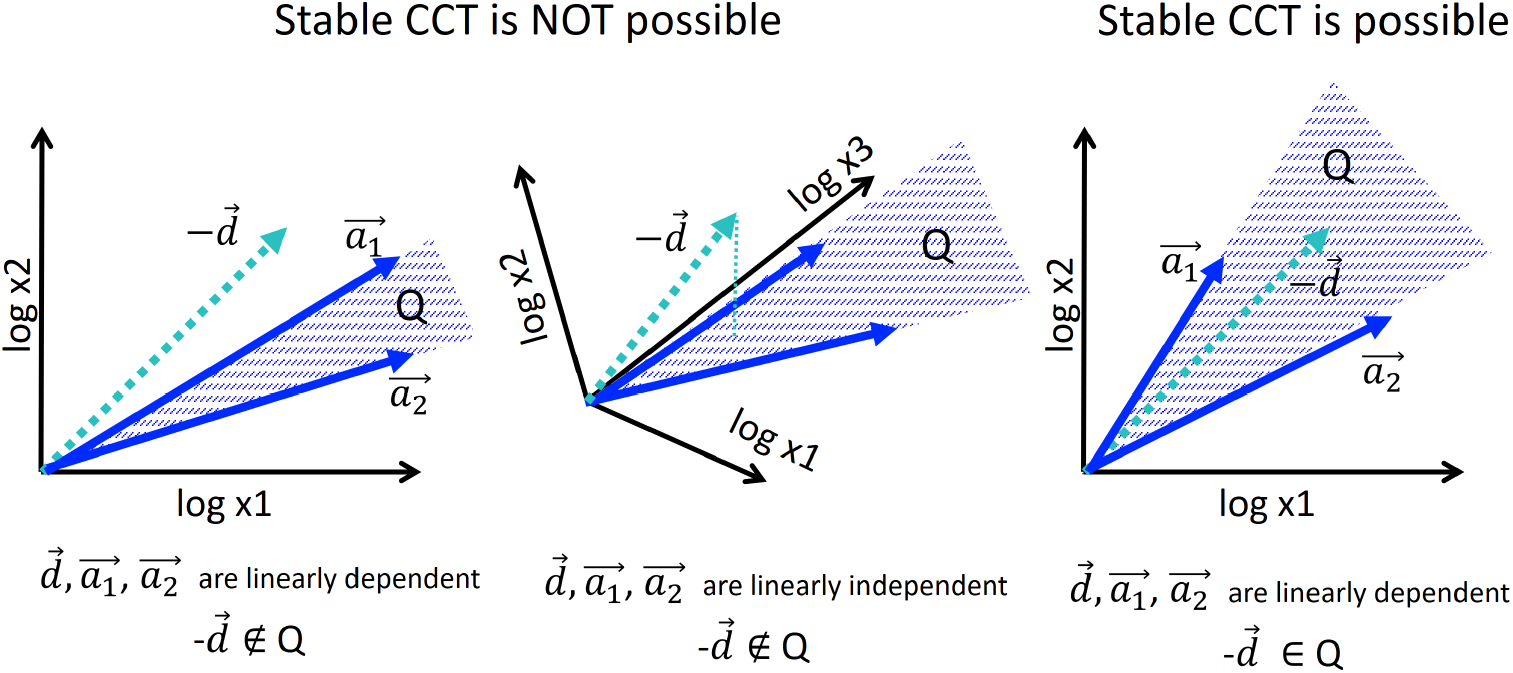
Condition of existence of stable cell cycle trajectory in the model of allometric growth with switches. For illustration, only two growth vectors *a*_1_, *a*_2_ are considered, and 2D or 3D embedding space. Stable piecewise linear trajectory is possible only if the negative of the cell division vector –*d* belongs to the convex cone 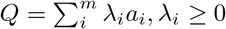. Only in this case, the cyclic equality 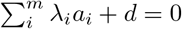 is possible. In general position, the condition can be assured only when *m* ≥ *n*, where n is the dimensionality of the trajectory space (see text for the formal proof).

Note that the proven Theorem is more general than the model of allometric growth with switches itself since it does not assume any particular shape of the switching surfaces *L_k_*: they can be linear or nonlinear. Another generality consists in that the vector *d* can have any non-zero coordinates, not necessarily equal to – log 2.

Examples in Figures 2, 3 correspond to the case *n* = 2, *m* > *n*. For example, the cell cycle trajectory modeled in Figure 3 contains *m* = 4 segments in 2D, which makes the vectors *a_i_* ∈ *R*^2^, *i* = 1…4 linearly dependent, and, of course, *d* ∈ *R*^2^. The cell cycle model based on allometric growth is not contradictory in this case.

Now let us formulate our second statement. We can recall that vectors *a_i_* are confined to the *n*-dimensional intrinsic subspace of CCT by projection from the multi-dimensional ambient space of all elementary measurements. Choice of *n* depends on our estimate of the CCT intrinsic dimesionality. However, movement along vectors *a_i_* can be also seen in the complete space with thousands of coordinates. In this space for sufficiently small *m*, any *m* vectors will almost always be linearly independent. Only projection into smaller than *m*-dimensional space will guarantee that these vectors are linearly dependent. This makes us hypothesize: if *m* segments are observed in CCT piecewise linear approximation in any linear projection then the most natural choice for *n* is at least *m*, i.e. *n* ≥ *m*. Combining two statements together (*m* ≥ *n* and *n* ≥ *m*) allows us to state that the correspondence *m* = *n* is the most natural expectation for a cell cycle trajectory.

We explicitly verified this correspondence for the trajectory shown in Figure 1. The curvature analysis suggests the existence of 5 segments for the cell cycle trajectory reconstructed in the subspace of 30 first principal components of the complete dataset. However, some of these components might correspond to the variance not related to the progression through the cell cycle. In order to diminish the possible role of this variance, we considered a reduced version of the dataset confined to cell cycle-related genes only. We estimated the global intrinsic dimensionality, using six different linear ID estimators from skdim Python package https://github.com/j-bac/scikit-dimension, and it varied from 2 to 7, with average value 4.0. The scree plot shows existence of two dominant eigenvalues explaining 83% of total variance, indicating that the trajectory is relatively flat and located close to a 2D linear manifold. However, the residual variance demonstrated visible patterns related to transcriptional epochs in at least the first four principal components (Figure 7). The distribution of projections on the first four principal components well separated some transcriptional epochs (Figure 7, diagonal). Also, projections in higher dimensions highlighted the existence of sharp turning points between the segments which were less clear in the 2D projection on the first two principal components.

In addition, we split the data points into 5 classes according to projection on 5 segments of the principal curve (0-A, A-B, B-C, C-D, D-E), each of which is approximately linear. For each of this class, we computed the unity vector corresponding to the direction of the first principal component in the space of cell cycle genes with 198 dimensions. Afterwards, we estimated the effective rank of the matrix composed of 5 vectors representing the directions of the transcriptional epochs in the multi-dimensional space (see Methods), and it appeared to be 4, which indicates to that at least 4 out of 5 vectors determining the trajectory segments can be considered linearly independent.

As a result, we concluded that the embedding dimensionality for the transcriptomic cell cycle trajectory can be estimated as close to four. Therefore, restricting the trajectory to the plane of aggregate collective expressions of genes associated with S phase and G2/M phase (which roughly corresponds to the first two principal components) is a useful but incomplete approximation of CCT dynamics. Our reasoning suggests searching for additional biologically meaningful and statistically independent variables describing the progression through the cell cycle. The concrete gene expression dynamics shown in Figure 1,B provides a hint in this direction, but a careful and complete investigation of this question should be a subject of a separate study.

### 2.6 Extending the modeling formalism to piecewise smooth trajectories: simple kinetic model of cell cycle at transcriptomic level

The piecewise-linear model of automaton with switches described in the previous sections is phenomenological and lacks any notion of physical time and connection to the underlying kinetics of the lumped expression of genes involved in S phase and G2/M phases. A simple way to make it more concrete but still analytically tractable consists in introducing explicit processes of synthesis and degradation of the corresponding quantities, with kinetic rates changing in time. The simplest form of such dependence is piecewise-constant, with changes in the value of kinetic rates corresponding to the observed switches between transcriptional epochs of cell cycle progression.

Assuming the same epochs of cell cycle progression as above, and the same notations for variables (*S, M*, lumped expression of genes involved in S and G2/M phases correspondingly), their dynamics can be expressed as:

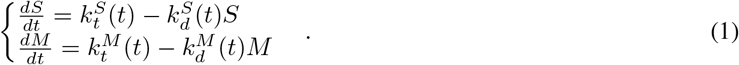

These equations must be accompanied by circular boundary conditions

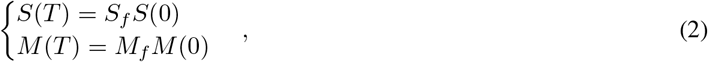

where *S_f_, M_f_* > 1 are some numbers describing the drop of the lumped cell cycle variables after the moment of cell division. The most natural choice for them is *S_f_, M_f_* = 2, as before: however, here we prefer not to fix these parameters and rather fit them from the actually observed trajectory.

There exist several reasons for which *S_f_* and *M_f_* might appear in the range 1 ≤ *S_f_, M_f_* ≤ 2 and not be equal. The most important of them is the technical biases introduced by sampling a limited amount of RNA, in the process of single cell transcriptome sequencing. It can lead to the situation when after cell division, the amount of RNA decreases non-uniformly between molecular processes. In particular, in all our experiments, we do observe the total amount of RNA reads does not decrease exactly by 2.0 and is rather close to 1.7-1.8. The decrease of the individual gene expression after cell division in terms of the number of reads, forms a bell-shaped distribution around this value with standard deviation close to 0.2.

The equations (1) with piecewise-constant in time kinetic rates and the boundary conditions (2) can be solved analytically for arbitrary number of levels in the piecewise-constant functions *k_t_*(*t*), *k_d_*(*t*). The resulting dynamics in the plane log *S*(*t*), log *M*(*t*) represents a cell cycle trajectory parametrized by physical time, which consists of piecewise-smooth segments of three types. If a segment is characterized by 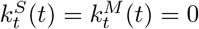 then the corresponding segment is linear in the logarithmic coordinates (since the underlying dynamics is exponentially decaying). If a segment is characterized by 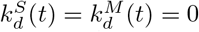 then the corresponding segment is also linear in both logarithmic and initial coordinates. For a segment where at least one degradation 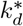 and one production kinetic rate 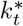 are positive, the dynamics follows a nonlinear curve in the logarithmic space, which remains monotonous (each of the coordinates does not change the derivative sign). The nonlinearity of the segment becomes important when one of the variables is at exponential increase or decrease stage, while the other is at linear or close to saturation stage. Otherwise, the segment remains to be close to a line in logarithmic coordinates.

In order to choose the number of constant levels of the kinetic rates, we studied the averaged RNA velocity values along the cell cycle as a function of pseudotime (see Figure 6,A,B). For the *S* variable, we decided to keep only one non-zero level of 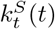 during the transcriptional epoch Ts, and two levels of 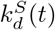, one for the exit from mitosis epoch and one for the rest of the dynamics. The choice was similar for *M* variable, but we took into account that a boost of expression of the lumped *G*2/*M* genes is visible in the beginning of the transcriptional epoch T2, just after switching off the S phase genes. During mitosis we assumed that all production rates are zero, corresponding to the lack of transcription in the M phase. The resulting choice of levels for the kinetic rates is shown in Figure 6,C.

**Figure 5:**
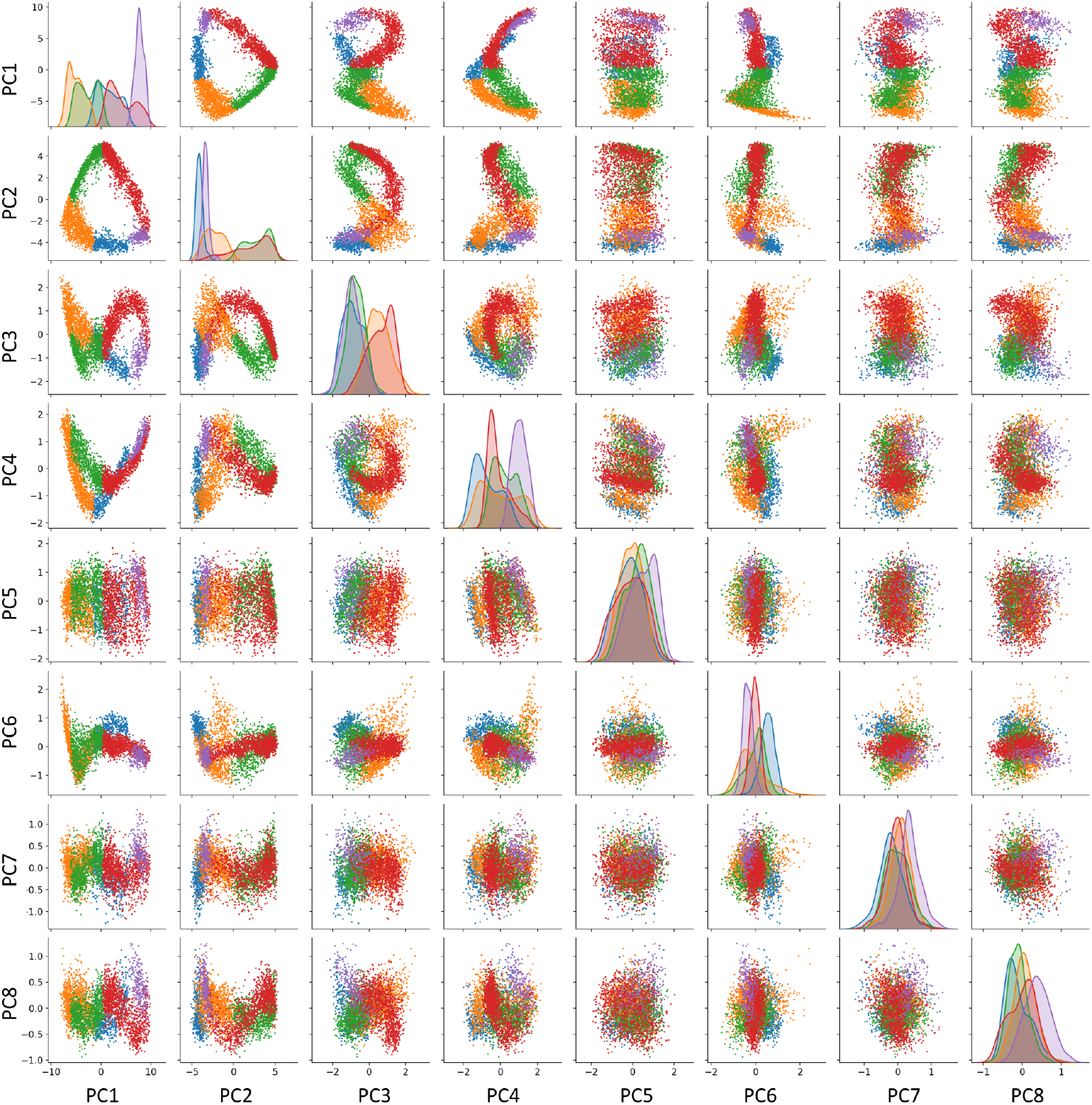
Visualizing the transcriptomic cell cycle trajectory of CHLA9 cell line in projections on first 8 principal components, computed in the subspace of known cell cycle genes. The data points are partitioned accordingly to the segmentation of the CCT into 5 transcriptomic epochs, also shown in Figure 1, 0-A (blue), A-B (orange), B-C (green), C-D (red), D-E (purple).

**Figure 6:**
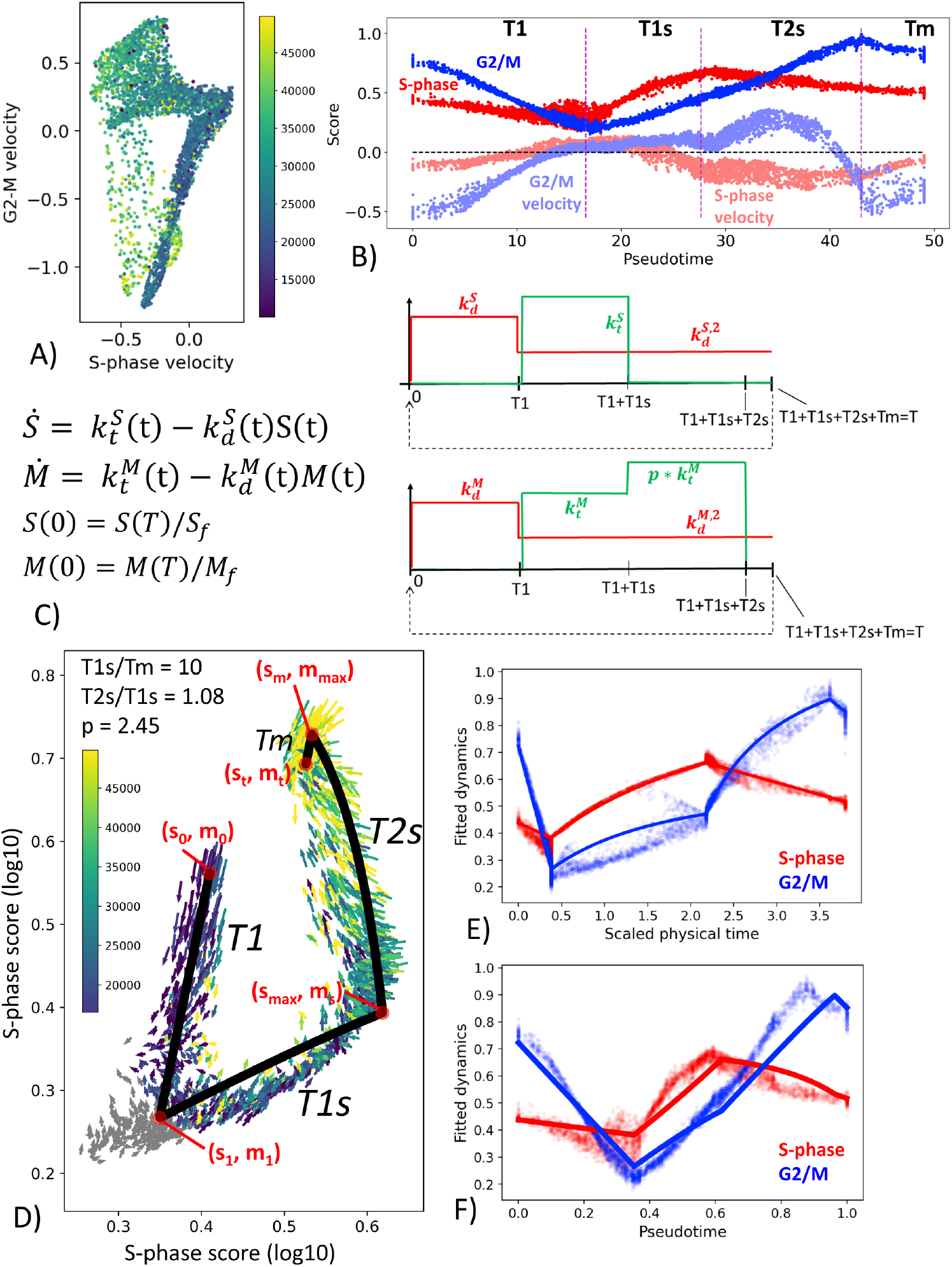
Simple kinetic model of cell cycle transcriptome dynamics. A) Mean RNA velocity values for S-phase and G2/M genes. B) Pseudotemporal dynamics of S-phase and G2/M scores (shown with more intense color) and mean RNA velocity values (shown with semi-transparent color). C) Description of the simple kinetic model of cell cycle transcriptome. Model equations are shown on the left and the changes in the values of kinetic rates (degradation, in red, and synthesis, in green). D) Result of fitting the model dynamics to cell cycle transcriptome dynamics observed in CHLA9 cell line. E),F) Inferred real-time and pseudotemporal dynamics of cell cycle transcriptome in CHLA9 cell line.

**Figure 7:**
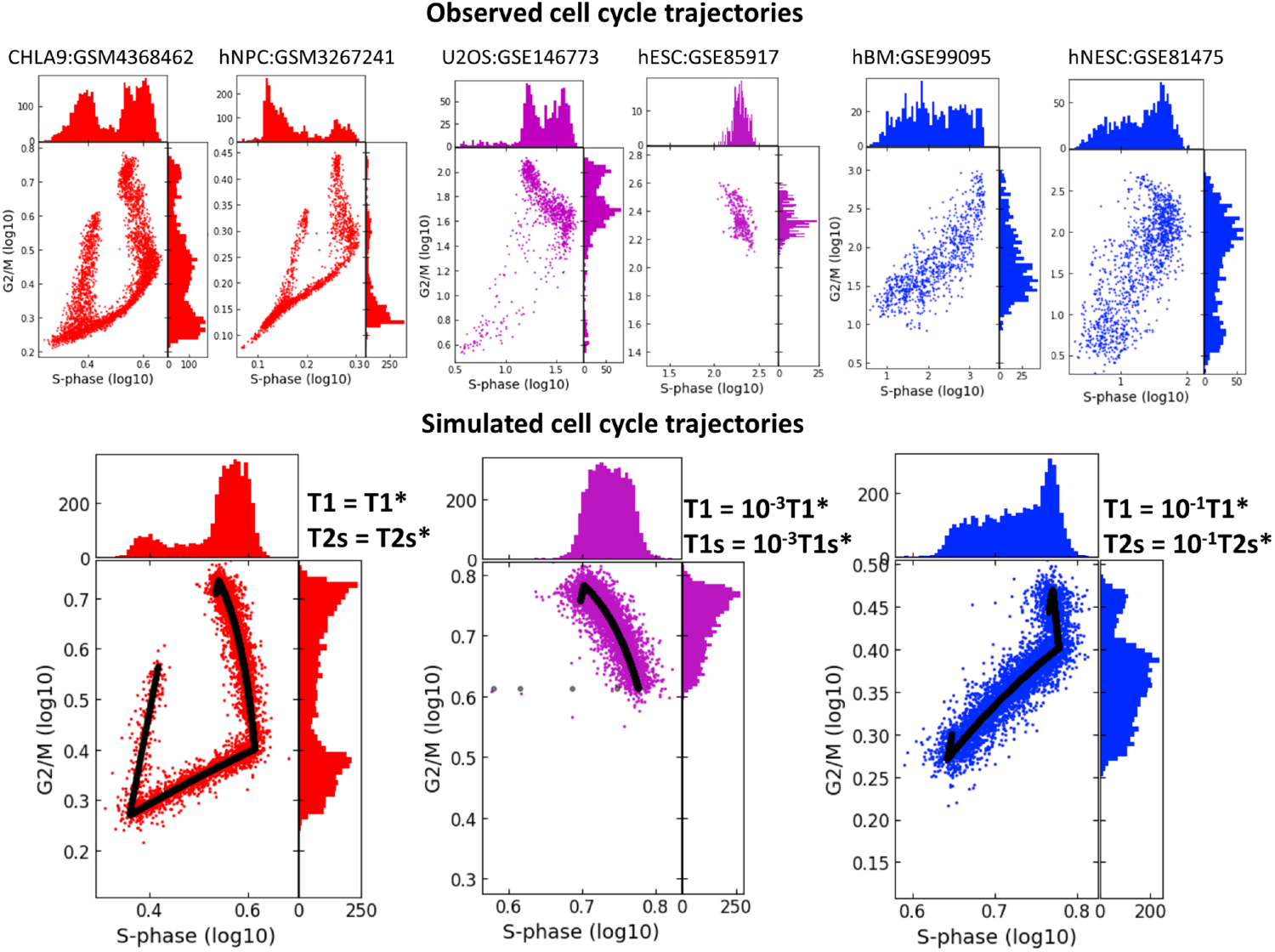
Studying the effect of shortening the durations of transcriptional epochs T1 and T1s or T1 and T2s on the geometry of cell cycle trajectory projected onto the S-phase and G2/M-phase scores plane. The simulated trajectories (in the lower part of the figure) are produced by taking the parameters of the CHLA9 fit of model dynamics (red plot) and changing the durations of T1 and T1s epochs (violet plot) or the durations of T1 and T2s epochs (blue plot). Each simulation shows the trajectory (black line) sampled with Laplacian noise added, with score distribution histograms shown at the plot margins. The upper part of the plot shows six real-life cell cycle trajectories observed in different systems, with GEO identifiers indicated. In each plot title either cell line name is provided, or hNPC means human neural precursor cells, hESC - human embryonic stem cell, hBM - human bone marrow, hNESC - human neural epithelial stem cell.

The advantage of the proposed simple model of cell cycle trajectory is in that it is fully analytically tractable and its parameters can be uniquely fit to the cell cycle trajectory observed from single cell data, given some biologically meaningful constraints. Thus, assuming that the duration of mitosis is by order of magnitude faster than the T1s epoch, for CHLA9 cell line one estimate the ratio between transcriptional epochs T2s and T1s as close to 1.0 and the value of transcriptional boost of G2/M genes in T2s epoch as close to 2.5-fold (Figure 6,C). The determined values of all other parameters can be found in the Jupyter notebook at https://github.com/auranic/CellCycleTrajectory_SegmentModel.

### 2.7 Simulating cell cycle trajectories with various durations of temporal transcriptional epochs

After fitting the kinetic parameters for an observable in the S-phase vs G2/M score plane cell cycle trajectory, one can perturb the parameters and investigate how the trajectory geometry depends on them.

In real life scRNASeq datasets, we observe that CCT geometry can appear very different in various biological systems. In projection onto the plane of standard scores of S-phase and G2/M phase genes, scRNASeq datasets might reveal or not the circular nature of CCT. In some cases, the circular structure is not at all detectable via this projection, see Figure 7, and the two scores might be connected via strong positive or negative correlation. Also, in some systems we observed co-existence of several CCT shapes, like it is the case in the U2OS cell line dataset (GSE146773). The univariate histograms of two score distributions might be characterized by bi- or uni-modal character.

Quite strikingly, we were able to reproduce these patterns qualitatively by fitting the kinetic parameters to the CHLA9 scRNASeq dataset, and then by manipulating the durations of *T*1, *T*1s and *T*2*s* transcriptional epochs and producing computer-simulated trajectory examples. Thus, significant reduction in the duration of both *T*1 and *T*1s epochs led to the negative correlation pattern between S-phase and G2/M scores. This could be interpreted as drastic reduction of the G1 cell cycle phase. In real life datasets, such pattern has been observed in human embryonic stem cells (dataset GSE85917).

If both *T*1 and *T*2*s* were shortened then this led to the increase of the positive correlation between two scores, Figure 7. This pattern was indeed observed in human bone marrow and human neural epithelial stem cell-related single cell datasets (GSE99095 and GSE81475).

### 2.8 Predicting cell line doubling time from the geometrical properties of cell cycle trajectory

The developed simple kinetic model leads to a simple prediction which can be validated: *the total length of the transcriptomic cell cycle trajectory must diminish in rapidly dividing cells*. This can be interpreted as a consequence of the fact that in rapid proliferation process, during the post-mitotic G1 phase (T1 transcriptional epoch) there is not enough time to degrade all mitotic transcripts produced before the cell division moment, so they are reused in the consequent cell cycle phases.

We verified this prediction in a relatively large collection of cell line scRNASeq datasets. Using the data from Cellosaurus database, we identified those few ones for which the cell line doubling time has been estimated, and for which the number of available single cell profiles passed the quality control exceeded 300 cells.

We used the total length of the principal circle fit in the 2D plane of the scaled to max one cell cycle phase scores, as a proxy to quantify the level of CCT contraction (see Methods). This measure was correlated with cell line doubling time in hours. Two cell lines CHLA10 and SCC25 appeared to be strong outliers from otherwise significant positive regression line (Pearson correlation 0.931, p-value=1θ^−5^), see Figure 8. When this regression line was used as a predictor, CHLA10 cell line was predicted to have doubling time around 64 hours (instead of determined by database search 32 hours) and for SCC25 around 78 instead of 50 hours. It is known that cell line doubling time can vary depending on the growth conditions, so we hypothezised that this variability could explain the appearance of two outliers. If two of them were kept in the regression calculation, it remained significant but less strong (Pearson correlation 0.67, p-value=0.01).

**Figure 8:**
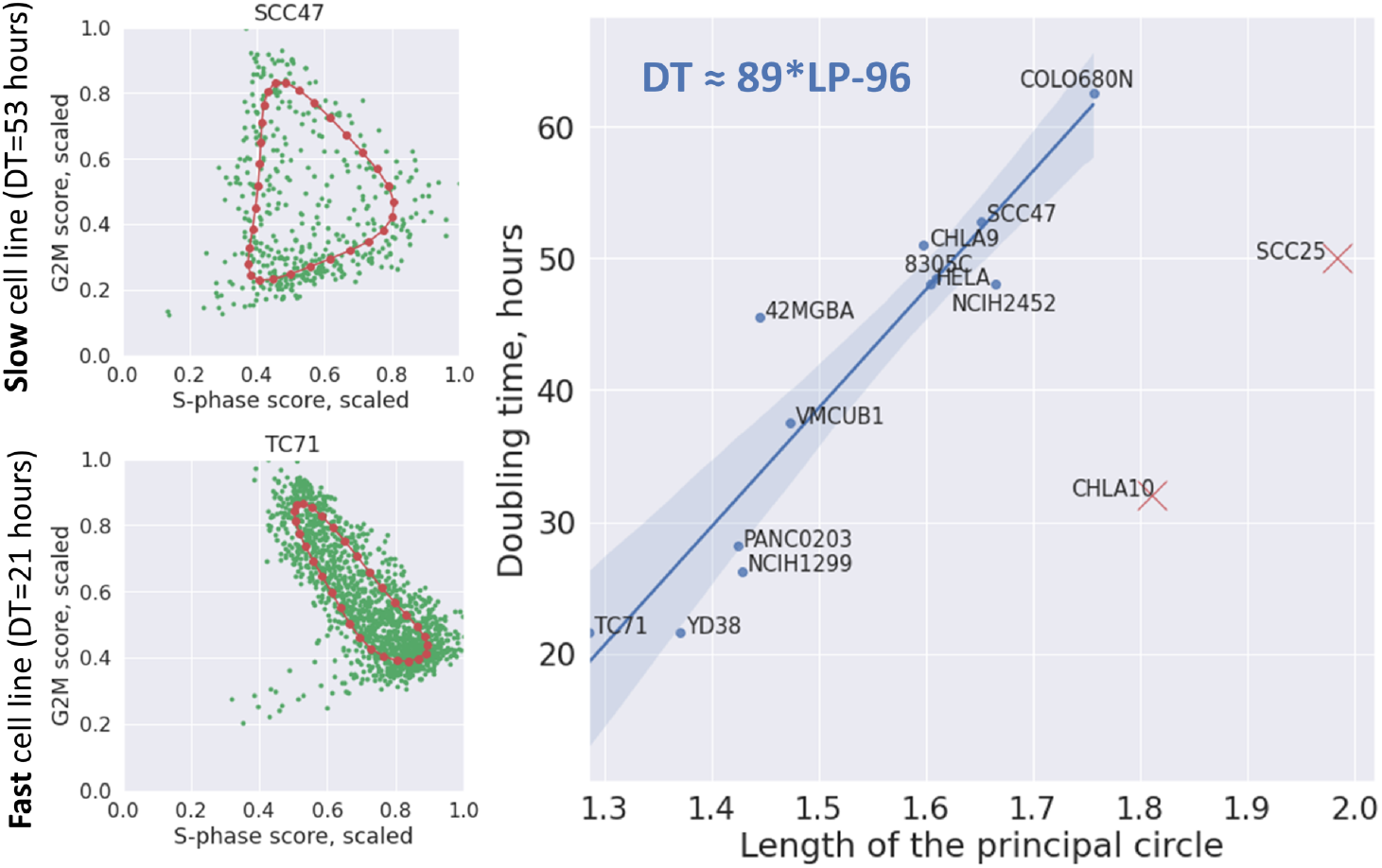
Dependence of cell line doubling time (DT) on the length of the principal circle approximating the cell cycle trajectory in the 2D plane of scaled (divided by the maximum value) S-phase and G2M scores. On the left two examples of principal circles are shown in red, and cells in green. On the right the linear regression line with confidence intervals is shown connecting the length of the principal circle with cell line doubling time (Pearson correlation 0.931, p-value=10^−5^). The regression formula is shown on the plot in top left corner. Two cell lines indicated by red crosses were eliminated from the regression as evident outliers.

## 3 Methods and materials

### 3.1 Single cell data used in this study

We made a screening of available single cell sequencing of cancer cell lines in order to identify datasets with sufficient number of good quality single cell transcriptomic profiles and in which the principal source of transcriptomic heterogeneity would be progression through the cell cycle. We identified publicly available scRNASeq data on CHLA9 Ewing sarcoma cell line, produced with 10x Genomics sequencing technology [26], which contained more than 4000 cells with total number of UMIs varying from 10000 to 50000, after applying the standard quality criteria and filtering cells containing a large fraction (>20%) of reads in mitochondrial genes. For this dataset, we reanalysed the raw sequencing data using Kallisto mapper [31] resulting in a loom file that could be used for obtaining the gene expression levels and for quantifying RNA velocity vectors [32].

In addition, we used a recently published collection of 200 scRNASeq profiles of cancer cell lines from Cancer Cell Line Encyclopedia (CCLE) collection [33]. We also analyzed several scRNASeq datasets by downloading them directly from Gene Expression omnibus (GEO).

The estimation of cell line doubling times, when available were obtained from Cellosaurus database [34]

### 3.2 Definition of cell cycle genes

We systematically tested several definitions of cell cycle genes and verified that the final results are not strongly influenced by the concrete choice of them. In our experiments we used the following definitions:

- Standard “Regev’s set”: markers of S- and G2/M cell cycle phases used in scanpy tutorials [35]
- Set of cell cycle genes annotated in Reactome pathway database [36]
- Set of top-contributing genes, extracted from application of independent component analysis (ICA) of the concrete dataset, from those components whose top-contributing genes were strongly associated with the cell cycle. In particular, similarly to our previous work [27], two independent components very significantly matched the markers of S- and G2/M cell cycle phases in all single cell cell line datasets we analyzed.

Cell cycle phase scores were computed as an average expression of marker genes for the corresponding cell cycle phase in log scale, which roughly corresponds to the geometric mean of the raw count measures.

### 3.3 Pooling reads from neighbouring cells for leveraging the technical drop-out

We found out that the analysis of CCT is significantly improved when standard pooling approach was applied to the raw count data, using an initial estimate of cell-to-cell proximity. More precisely, we used the standard initial standard data normalization and dimensionality reduction in order to compute the distances between cells and construct the initial kNN graph, which was used to pool row reads from a cell and all its *k* nearest neighbours. In our experiments we used *k* = 10 and *n* = 30 components for reducing the data dimensionality during normalization. Pooled read counts were used for final normalization, but the initial total read counts per cell measure were kept for visualization and further analysis.

### 3.4 Cell cycle trajectory-based single cell data normalization

Total number of reads in a cell represents a strong signal in proliferating cell populations. By itself it is an extensive value such that it should be divided (approximately) by half in the process of cell division. In our modeling we needed a description of the cell state in terms of extensive values of gene expression levels measured such that they would be also divided approximately by two on average after the moment of cell division. Therefore, the widely used global library size normalization did not suit our purposes, since after global library size normalization, cell division does not lead to halving the total number of reads.

At the same time we observed that without any library size normalization, the cells presumably located at similar stages of cell cycle progression could be characterized by a wide range of total number of reads, probably caused by technical variability factors. Therefore, library size normalization was required but not at the global cell population level. We hypothesized that the total number of reads should increase in the course of cell cycle progression on average such that the cells characterized by similar value of pseudotime along the cell cycle trajectory could be normalized to the same local library size. As usual, this poses a chicken-or-egg problem because for reconstructing the cell cycle trajectory one needs normalized data, and for normalization of the library size one needs a reconstructed trajectory. This problem is similar to those approaches which use normalization locally conditioned on clusters in single cell datasets [37].

We used a simplified two-stage approach for library size normalization which preserved both the geometric structure of CCT and the trend of increasing the total number of reads along CCT.

1. The row count data have been normalized to the global median number of counts and ln(x+1)-transformed, using standard functions of scanpy. 10000 most variable genes have been selected, the dimensionaity was reduced to 30 by PCA. In the reduced space, a kNN graph has been computed using the standard Euclidean distance for k=10. This graph was used for pooling reads from neighbour cells as described above.
2. For such initially normalized dataset, we computed closed cell cycle trajectory in the subspace of cell cycle genes, by fitting a principal closed curve, using the Python implementation of ElPiGraph [38]. The data points were partitioned according to the proximity to the nodes of the elastic principal curve.
3. In each partition we analyzed the distribution of the total number of reads across cells. We performed correction of cell-to-node assignment by splitting an anomalously wide partition between two neighbouring partitions. The anomalously wide partition corresponded to the moment of cell division since it contained both cells at the very end of cell cycle progression with the largest number of reads and cells just after cell division event containing the minimal number of reads. Splitting this distribution allowed us to distinguish cells just before and just after the cell division into distinct partitions.
4. The median total number of counts in each resulting corrected partition was computed. The median values of the total number of reads in the cells of each partition have been smoothed by univariate spline or a piecewise-linear function of pseudotime, taking into account the cyclic boundaries of the trajectory.
5. Each cell’s library size was normalized to the smoothed local median value of the total number of reads.
6. The newly normalized pseudocount data matrix passed through the same pre-processing as described in 1), namely a) Pooling reads from neighbour cells using the kNN graph obtained with trajectory-based normalized data, b) ln(x+1) transformation, selecting most variable 10000 genes.

The cell cycle trajectory-based normalization procedure is illustrated in the Jupyter notebook at https://github.com/auranic/CellCycleTrajectory_SegmentModel, which can be easily reused for other cell lines.

### 3.5 Computing the cell cycle trajectory and quantifying pseudotime

We used the ElPiGraph Python package to fit elastic principal curves or closed elastic principal curves (principal circles) to single cell data distributions [38]. ElPiGraph was applied in the data space defined by the set of 10000 most variable genes or by the cell cycle-related genes, after dimensionality reduction by PCA (first 30 principal components were retained). In order to compute open elastic principal curve with *q* nodes, first a closed curve was fit with *q*/2 nodes, then a node with the least number of data points projected into it was removed from the principal graph, and this configuration was used as initialization to compute the elastic principal graph without branching having *q* nodes.

The pseudotime *s_i_* for a data point *x_i_* was computed as a continuous geodesic distance measured from the root node to the projection of *x_i_* onto the principal curve, quantified in the units of the number of edges. Therefore, the value of the pseudotime was in the range [0, *q* – 1], where *q* is the number of nodes. The root of the principal curve was chosen as one of its ends, such that the value of the initial total number of reads would increase as a function of pseudotime.

### 3.6 Curvature analysis of the cell cycle trajectory

In order to compute the Riemannian curvature of the principal curve defined by the position of its nodes in the multidimensional space *y_i_* ∈ *R^n^, i* = 1… *q*, the node coordinates were first represented as *n* functions of the natural parameter (pseudotime) *s*, 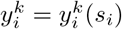, *i* = 1…*q, k* = 1…*n*. The value *s_i_* for each node was taken as anumber of edges of the elastic principal curve connecting the node *i* to the root node. Each set of numbers 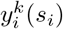, *i* = 1…*q* was interpolated by a cubic univariate spline *y^k^*(*s*). In each node *i* of the curve the curvature was evaluated as 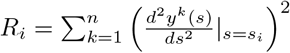.

### 3.7 Estimating the effective dimensionality of a set of vectors

In order to estimate the effective dimensionality of CCT, we used skdim Python package available at https://github.com/j-bac/scikit-dimension. We used linear estimators of global intrinsic dimensionality, based on application of PCA and various approaches to select the significant number of eigenvalues from the scree plot.

In order to compute the effective rank of a rectangular matrix, we looked at the distribution of its singular values, and selected such a number of them that the ratio between the largest and the smallest number would not exceed 10, such that the reduced matrix is well-conditioned.

### 3.8 Fitting parameters of the simple kinetic cell cycle model

Using the choice of levels for piecewise constant kinetic rates shown in Figure 6,C, we could derive the dependence of the initial state of the cell cycle from the kinetic rates and the durations of four transcriptional epochs *T*_1_, *T*_1*s*_, *T*_2*s*_, *T_m_*:

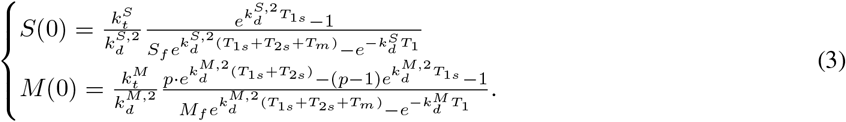

Starting from the initial point of the trajectory *S*(0), *F*(0) it is possible to analytically write down the coordinates of all other borders of the transcriptional epochs:

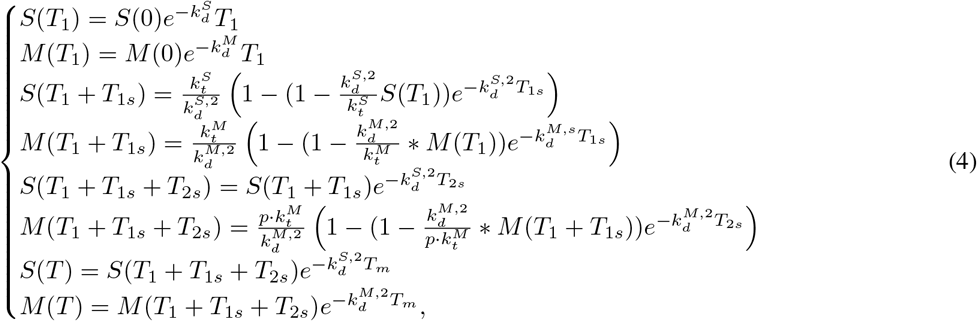

where *T* = *T*_1_ + *T*_1*s*_ + *T*_2*s*_ + *T_m_* is the full duration of the cell cycle. One can estimate the position of these points from the analysis of observed cell cycle trajectory curvature ((*s*_0_, *m*_0_), (*s*_1_, *m*_1_), (*s_max_, m_s_*), (*s_m_, m_max_*), (*s_t_, m_t_*), shown by red points in Figure 6,D)) by requiring that the model trajectory should pass as close as possible to them. This defines an optimization problem which can be easily solved numerically by iterations, using the simplest fixed-point algorithm. The details of parameter fitting are provided in the Jupyter notebook at https://github.com/auranic/CellCycleTrajectory_SegmentModel.

We note that this optimization does not allow us to determine all the model parameters uniquely, since they enter in the aforementioned optimization functional as certain combinations (as simple rational functions), namely, 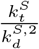, 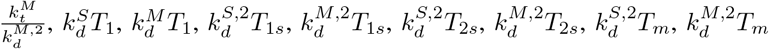. Two other parameters *M_f_, S_f_* define the observed cell division vector in (2). One extra parameter p denotes transcriptional production acceleration of *G*2/*M* genes during the transcriptional epoch *T*_2*s*_ compared to the transcriptional epoch *T*_1*s*_ (Figure 6,C). Not all these quantities are independent, some of them are connected through nonlinear relations:

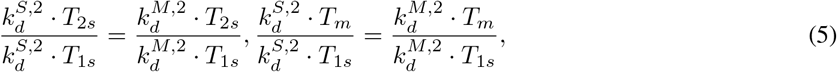

which overall gives 11 independent combinations of parameters provided 10 measurable coordinates of cell trajectory turning points in Figure 6,D.

Altogether, this means that 1) one needs to introduce at least one additional constraint in order to make the trajectory reconstruction unique and 2) physical time of the epochs *T*_1_, *T*_1*s*_, *T*_2*s*_, *T_m_* can not be uniquely computed from the cell cycle trajectory observed in the plane of S-, G2/M-phase scores. From the analysis of equations (5) it follows that the model can be uniquely parametrized if one will constraint one of the three quantities *p*, 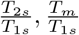. Finally, it is convenient to fix the durations *T*_1_, *T*_1*s*_ to some arbitrary values which allows to determine parameters 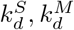 and the ratios 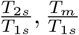.

In our numerical experiments we fixed the values of *T*_1_ and *T*_1*s*_ to their corresponding pseudotemporal durations (as the corresponding fractions of the total length of the cell cycle trajectory). We also fixed the ratio 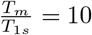, assuming that the mitosis must be fast in physical time compared to the transcriptional epoch including activating the expression of the genes involved in the S-phase.

### 3.9 Code availability

The Python notebooks allowing the reader to reproduce all the computations presented in this manuscript are freely available from https://github.com/auranic/CellCycleTrajectory_SegmentModel.

## 4 Discussion

This paper provides an analytical framework for treating the properties of cell cycle trajectory observed in single cell populations. Unlike the previously suggested model of the trajectory as a flat circle, we provide arguments that at least in some conditions the piecewise-linear in logarithmic coordinates approximation appears to fit the single cell transcriptomic data and to be biologically tractable. In particular, it allows us to delineate transcriptional epochs of cell cycle at which the corresponding segment of the trajectory remains close to linear in logarithmic coordinates which corresponds to locally allometric changes of the transcriptome.

We suggest two modeling formalisms to recapitulate the cell cycle transcriptomic dynamics as a sequence of switches. The first one is purely phenomenological and describes the dynamics as a change of states of a hidden automaton, leading to the switches of parameters of allometric growth, followed by a shift representing the cell division event. The advantage of this formalism is that it allows us to treat most general properties of cell cycle trajectory geometry.

In particular, we could prove a fundamental theorem on the number of intrinsic cell cycle states, which connects the number of linear segments in the trajectory and the embedding dimensionality of the cell cycle trajectory. The nature of this theorem, relying on “general position”-type arguments, is reminiscent of the well-known results imposing constraints on the number of the system’s internal states and the effective dimensionality of its environment, in several fields of science. For example, the Gause’s law of competitive exclusion and its generalizations states that the number of competing species is limited by the effective number of resources, characterizing the environment [39, 40]. The famous Gibbs’ phase rule in thermodynamics connects the effective number of the intensive variables with the number of components and phases in a system at thermodynamic equilibrium [41, 42]. All these results are also similar in terms of practical difficulties related to determining the effective system’s dimensionality.

From the physico-chemical point of view, the effective dimensionality is the number of the substances “lumps” in the cell cycle kinetics. Lumping-analysis produces a partition of all chemical species into a few groups and then considers these groups (“lumps”) as independent entities [43]. “Amounts” of these lumps are the combinations of the amounts of the chemical species [44, 45]. Theorem on the number of intrinsic cell cycle states means that the number of lumps n does not exceed the number of the internal states of the cell cycle transcription machinery. This means that kinetics allows reduction of the huge-dimensional space of all components to *n* ≤ *m* number of aggregated lumps.

The second modeling formalism that we suggested connects the geometric properties of the cell cycle trajectory to the underlying transcriptional kinetics and physical time. It uses the simplest chemical kinetics equations with kinetic rates represented as piecewise-constant functions of time. We show that the suggested model is fully analytically tractable and, under some biologically transparent assumptions, allows unique determination of its independent parameter combinations. This type of modeling allowed us to explicitly study the relation between pseudotime and physical time.

The precise connection between physical time and pseudotime (geometric time) in the cell cycle is worth studying in more detail since this is the central question in the dynamic phenotyping approach in general [46]. Some of these relations can be potentially quantified from exploring the variations of point density along the inferred trajectories [47]. Related to this, one can expect non-trivial phenomena in studying the cell cycle trajectory, such as effects of partial cell population synchronization under assumption of equal cell cycle durations in individual cells. This effect can lead to the appearance of density peaks in the reconstructed cell cycle trajectories that can not be explained by nonlinear relation between physical time and pseudotime [40].

As one of the applications of the suggested modeling formalism, we performed several numerical experiments on changing the durations of the transcriptional epochs overlapping with G1 or G2 cell cycle phases. We observed that these parameters might have a drastic effect on the shape of the CCT geometry and the form of the univariate variable distributions. This model prediction can be qualitatively confirmed by observing CCT properties of several in vitro and in vivo systems. The effect of CCT shrinkage might be relevant in characterizing the cell cycle properties in various conditions: for example, when one can manipulate the activity of an oncogene [27]. We show that the CCT geometry can be predictive to estimate the cell line doubling time which can be a proxy of cell cycle duration.

The relation between transcriptomic dynamics and the established definitions of cell cycle phases has been discussed and even quantified using standard molecular biology techniques [1, 13]. In this study, we deliberately leave open the question on defining the exact cell cycle phase borders from the transcriptomic CCT geometry. We found that this relation can not be the exact match: one of the reasons for this is delayed production of proteins, and dependence of the cell cycle progression from post-translational protein modifications. The transcriptomic dynamics is relatively slow, and activation of protein synthesis is switched on in advance, leaving time for producing enough proteins needed at a certain stage of the cell cycle molecular program. Same is true for the process of degradation of RNAs involved in cell cycle: a cell needs enough time after mitosis to degrade all cell cycle-related transcripts.

The suggested formalism is not limited to transcriptomic data. It looks promising to analyse the geometrical properties of cell cycle trajectory measured in unsynchronized cell populations profiled at various levels of molecular description, including epigenetics and protein expression, when the datasets of sufficient volume and quality will become available.

## 4.1 Acknowledgements

This work was supported by the French government under management of Agence Nationale de la Recherche as part of the”Investissements d’avenir” program, reference ANR-19-P3IA-0001 (PRAIRIE 3IA Institute), The work was supported by the Ministry of Science and Higher Education of the Russian Federation (Project No. 075-15-2021-634) and by European Union’s Horizon 2020 program (grant No. 826121, iPC project).

